# Optimizing EEG source reconstruction with concurrent fMRI-derived spatial priors

**DOI:** 10.1101/2021.02.19.431976

**Authors:** Rodolfo Abreu, Júlia F. Soares, Sónia Batista, Ana Cláudia Lima, Lívia Sousa, Miguel Castelo-Branco, João Valente Duarte

**Affiliations:** Coimbra Institute for Biomedical Imaging and Translational Research (CIBIT), Institute for Nuclear Sciences Applied to Health (ICNAS), University of Coimbra, Coimbra, Portugal; Faculty of Medicine, University of Coimbra, Coimbra, Portugal; Neurology Department, Centro Hospitalar e Universitário de Coimbra, Coimbra, Portugal

## Abstract

Reconstructing EEG sources involves a complex pipeline, with the inverse problem being the most challenging. Multiple inversion algorithms are being continuously developed, aiming to tackle the non-uniqueness of this problem, which has been shown to be partially circumvented by also including fMRI-derived spatial priors in the inverse models. However, only task activation maps and resting-state networks (RSNs) have been explored so far, overlooking the recent, but already accepted, notion that brain networks exhibit dynamic functional connectivity (dFC) fluctuations. Moreover, there is no consensus regarding the inversion algorithm of choice, nor a systematic comparison between different sets of spatial priors. Using simultaneous EEG-fMRI data, here we compared four different inversion algorithms (MN, LORETA, EBB and MSP) under a Bayesian framework, each with three different sets of priors consisting of: 1) those specific to the algorithm (S1); 2) S_1_ plus fMRI task activation maps and RSNs (S_2_); and 3) S_2_ plus network modules of task-related dFC states estimated from the dFC fluctuations (S_3_). The quality of the reconstructed EEG sources was quantified in terms of model-based metrics, namely the free-energy and variance explained of the inversion models, and the overlap/proportion of brain regions known to be involved in the visual perception tasks that the participants were submitted to, and RSN templates, with/within EEG source components. Model-based metrics suggested that model parsimony is preferred, with the combination MSP+S_1_ exhibiting the best performance. However, optimal overlap/proportion values were found using EBB+S_2_ or MSP+S_3_, respectively, indicating that fMRI spatial priors, including dFC state modules, are crucial for the EEG source components to reflect neuronal activity of interest. Our results pave the way towards a more informative selection of the optimal EEG source reconstruction approach, which may be crucial in future studies.

## 1 Introduction

Electroencephalography (EEG) measures the electrical potential differences between electrodes placed at different scalp sites that are generated by an ensemble of brain cells acting in synchrony. Because of its fairly direct relationship with neuronal activity and its remarkable temporal resolution at the sub-millisecond scale, EEG has proven pivotal for studying both healthy and abnormal human brain function in general, and particularly brain functional connectivity and its dynamics (Niedermeyer and Lopes Da Silva, 2005). However, the spatial identification and characterization of the brain networks underlying the electrical potentials measured at the scalp are not possible from these scalp signals alone, given the poor spatial resolution of the EEG at the centimeter scale (Michel et al., 2004). Fortunately, the continuous technological advances of EEG hardware and signal processing techniques now permit a reliable reconstruction of those brain networks (Abreu et al., 2020b), by localizing and estimating the strength of the neural generators responsible for the scalp EEG signals – the so-called EEG source reconstruction (Michel and Murray, 2012).

Reconstructing EEG sources involves a complex pipeline that can be divided into the forward and the inverse problems. The forward problem relates with the estimation of the impact of a given source in the brain on the scalp electrical potentials, and is typically solved by building realistic and subject-specific head models from individual structural magnetic resonance images (MRI) using well-defined processing pipelines. By incorporating the 3D localization of the scalp electrodes on these head models, a lead field can then be computed, establishing the relationship between the activity of the different sources in the brain and the signal measured at each electrode (Michel and Brunet, 2019). Conversely, the inverse problem relates with determining the sources in the brain that generate a given scalp distribution of electrical potential differences (i.e, EEG topography). Because of the non-uniqueness of its solution, the inverse problem is considered the most challenging, with a plethora of inversion algorithms being available for solving it (Michel et al., 2004). These can be roughly divided into current source density (CSD) estimates and beamformers (Grech et al., 2008; He et al., 2018). Despite the choice between the two types being application-dependent to some extent (Halder et al., 2019), CSD-based algorithms are the most commonly used in the literature; within these, the more recent distributed source localization algorithms are preferable over dipole source localization algorithms, as the latter require prior knowledge regarding the number of sources to be estimated. Several distributed source localization algorithms have been developed, the most common being the minimum norm (MN) solutions (Hämäläinen and Ilmoniemi, 1994) and their variations: weighted minimum norm (WMN; De Peralta-Menendez and Gonzalez-Andino, 1998), low resolution electromagnetic tomography (LORETA; Pascual-Marqui et al., 1994), local autoregressive average (LAURA; De Peralta-Menendez et al., 2004), among others (Michel and Brunet, 2019). However, there is no consensus regarding which algorithm yields the most accurate source reconstructions, and only a few studies have dedicated to systematically compare their performance. Motivated by the challenging task of defining a ground truth, most of these studies used simulations and reported inconclusive results, with the selection of the best algorithm highly depending on the simulated conditions (Bradley et al., 2016; Grova et al., 2006; Halder et al., 2019; Yao and Dewald, 2005). Similarly, the optimal inversion algorithm for reconstructing real EEG data has not been found yet (Hedrich et al., 2017); moreover, the associated results have not been appropriately validated based on the brain activity of interest, but rather using unspecific measures, typically the localization error, spatial spread and percentage of false positives, as well as the free energy and variance explained of the inversion model used when considering Bayesian frameworks (Michel and Brunet, 2019). Importantly, no study has focused on determining the extent at which the effects of different source reconstruction algorithms differ between groups in clinical studies. This is especially relevant in task-related fMRI studies, which are rapidly increasing in clinical research (Marinazzo et al., 2019).

In order to tackle the non-uniqueness of the inverse problem, assumptions and constraints need to be considered, which are reflected differently on each inversion algorithm. Their incorporation can be performed using two different approaches: by imposing penalty functions (Valdés-Sosa et al., 2009), or using a Bayesian framework (Friston et al., 2008; Trujillo-Barreto et al., 2004), particularly the parametric empirical Bayesian (PEB; Henson et al., 2010; Phillips et al., 2005). Although less popular, the PEB framework allows to describe a given assumption or constraint explicitly through appropriate postulated prior distributions, which can range from one as in the MN solutions (the identity matrix) to hundreds as in the multiple sparse priors (MSP) algorithm (Friston et al., 2008). This framework is thus extremely flexible for incorporating additional priors obtained from other imaging modalities, which has proved to be crucial for more efficiently circumventing the non-uniqueness of the inverse problem (Lei et al., 2015). The first studies used brain activation maps obtained from the analyses of task-based functional MRI (fMRI) data (Henson et al., 2010; Lei et al., 2012, 2011, 2010). More recently, the well-known brain networks that emerge from temporally correlated spontaneous fluctuations in the blood-oxygen-level-dependent (BOLD) fMRI signal (the so-called resting-state networks, RSNs) have also been used as spatial priors (Lei, 2012). In these studies, the spatial priors were derived from separately acquired fMRI data, which may scale down their potential for guiding the reconstruction of EEG, especially when focusing on spontaneous activity (Abreu et al., 2018). Additionally, task-based and resting-state functional networks are now known to continuously reorganize in response to both internal and external stimuli at multiple time-scales, resulting in temporal fluctuations of their connectivity – the so-called dynamic functional connectivity (dFC) (Hutchison et al., 2013). From dFC fluctuations, a limited, but variable, number of dFC states have been recurrently identified in the literature as the building blocks of brain functional connectivity (dynamics) (Preti et al., 2017), which are hypothesized to be associated with different cognitive, vigilance or pathological brain states (Thompson, 2018); however, they have not been considered as potential spatial priors for EEG source reconstruction so far.

Given the increasing relevance of EEG as a brain imaging tool, accurately estimating the underlying brain sources is critical in the study of both healthy and clinical populations. Considering the present limitations described in this section, here we compared four different inversion algorithms (MN, LORETA, empirical Bayes beamformer, EBB; and MSP), each with two different sets of additional fMRI-derived spatial priors (activation maps and RSNs, with and without including dFC states) on EEG data collected concurrently with fMRI at 3T from 6 multiple sclerosis (MS) patients and 7 healthy subjects performing visual perception tasks and during rest. The quality of the reconstructions was quantified through the free-energy and variance explained of the associated models, and in terms of the overlap between EEG source components and brain regions of interest associated with the tasks and RSN templates.

## 2 Materials and Methods

### 2.1 Participants

Six MS patients (mean age: 30±8 years; 2 males) and seven demographically matched healthy subjects (mean age: 30±6 years; 3 males) were recruited. The patients were selected by the clinical team at the Neurology Department of the University Hospital of Coimbra, and met the criteria for MS diagnosis according to McDonald Criteria (Thompson et al., 2018). All participants had normal or corrected-to-normal vision. The study was approved by the Ethics Commission of the Faculty of Medicine of the University of Coimbra, and was conducted in accordance with the declaration of Helsinki. All participants provided written informed consent to participate in the study.

### 2.2 Experimental protocol

The imaging session was performed at the Portuguese Brain Imaging Network (Coimbra, Portugal) and consisted of four functional runs: first, a functional localizer of the human middle temporal area (hMT+/V5, a low level visual area well-known to respond to simple motion patterns), followed by two runs of biological motion (BM) perception, and one final resting-state run.

The localizer run consisted of 10 blocks of 18 seconds, with each block comprising three periods: the first was a fixation period marked by a red cross positioned at the center of the screen for 6 seconds. During the second period, a 6 second pattern of stationary dots was shown, followed by the third (and final) period during which the dots were moving towards and away from a central fixation cross at a constant speed (5 deg/sec) for 6 seconds.

Biological motion stimuli were built based on human motion capture data collected at 60 Hz, comprising 12 point-lights placed at the main joints of a male walker. Each BM perception run consisted of 12 blocks of 40 seconds: 4 or 5 blocks (depending on the starting block) of the point-light walker facing rightwards or leftwards (*body* blocks), 4 or 5 blocks showing only the point-light located at the right ankle and moving rightwards of leftwards (*foot* blocks), and 3 blocks of the original 12 point-lights randomly positioned across the *y* axis, while maintaining their true trajectory across the *x* axis (*random* blocks). A total of 9 *body*, 9 *foot* and 6 *random* blocks were then collected during the two BM perception runs.

During the resting-state run, the participants were instructed to relax and only fixate a red cross positioned at the center of the screen.

### 2.3 EEG-fMRI data acquisition

Imaging was performed on a 3T Siemens MAGNETOM Prisma Fit MRI scanner (Siemens, Erlangen) using a 64-channel RF receive coil. In order to minimize head motion and scanner noise related discomfort, foam cushions and earplugs were used, respectively. The functional images were acquired using a 2D simultaneous multi-slice (SMS) gradient-echo echo-planar imaging (GE-EPI) sequence (6× SMS and 2× in-plane GRAPPA accelerations), with the following parameters: TR/TE = 1000/37 ms, voxel size = 2.0×2.0×2.0 mm^3^, 72 axial slices (whole-brain coverage), FOV = 200×200 mm^2^, FA = 68°, and phase encoding in the anterior-posterior direction. The start of each trial was synchronized with the acquisition of the functional images. A short EPI acquisition (10 volumes) with reversed phase encoding direction (posterior-anterior) was also performed prior to each fMRI run, for image distortion correction. Whole-brain, 1 mm isotropic structural images were acquired using a T_1_-weighted 3D gradient-echo MP2RAGE sequence.

The EEG signal was recorded using the MR-compatible 64-channel NeuroScan SynAmps2 system and the Maglink^™^ software, with a cap containing 64 Ag/AgCl non-magnetic electrodes positioned according to the 10/10 coordinate system, a dedicated electrode for referencing placed close to the Cz position, and two electrodes placed on the back for electrocardiogram (EKG) recording. Electrode impedances were kept below 25 kΩ. EEG, EKG and fMRI data were acquired simultaneously in a continuous way, and synchronized by means of a Syncbox (NordicNeuroLab, USA) device. EEG and EKG signals were recorded at a sampling rate of 10 kHz, synchronized with the scanner’s 10 MHz clock. No filters were applied during the recordings. The helium cooling system was not turned off, as it may carry the associated risk of helium boil-off in certain systems (Mullinger et al., 2008), and thus is not permitted in some clinical sites as the one used in this study. Respiratory traces were recorded at 50 Hz with a respiratory cushion from the physiological monitoring unit of the MRI system.

For each participant, 192 fMRI volumes were acquired during the localizer run, yielding 3.20 minutes of duration. The two BM runs had approximately 8.37 minutes, thus comprising 507 volumes each. The final resting-state run had approximately 8.08 minutes, corresponding to 485 volumes.

### 2.4 MRI data analysis

The main steps of the processing pipeline for deriving fMRI spatial priors (described here) and subsequently use them in EEG source reconstruction (described in the next section), as well as the metrics proposed for quantifying the quality of the source reconstruction, are depicted in Fig. 1.

**Figure 1.**
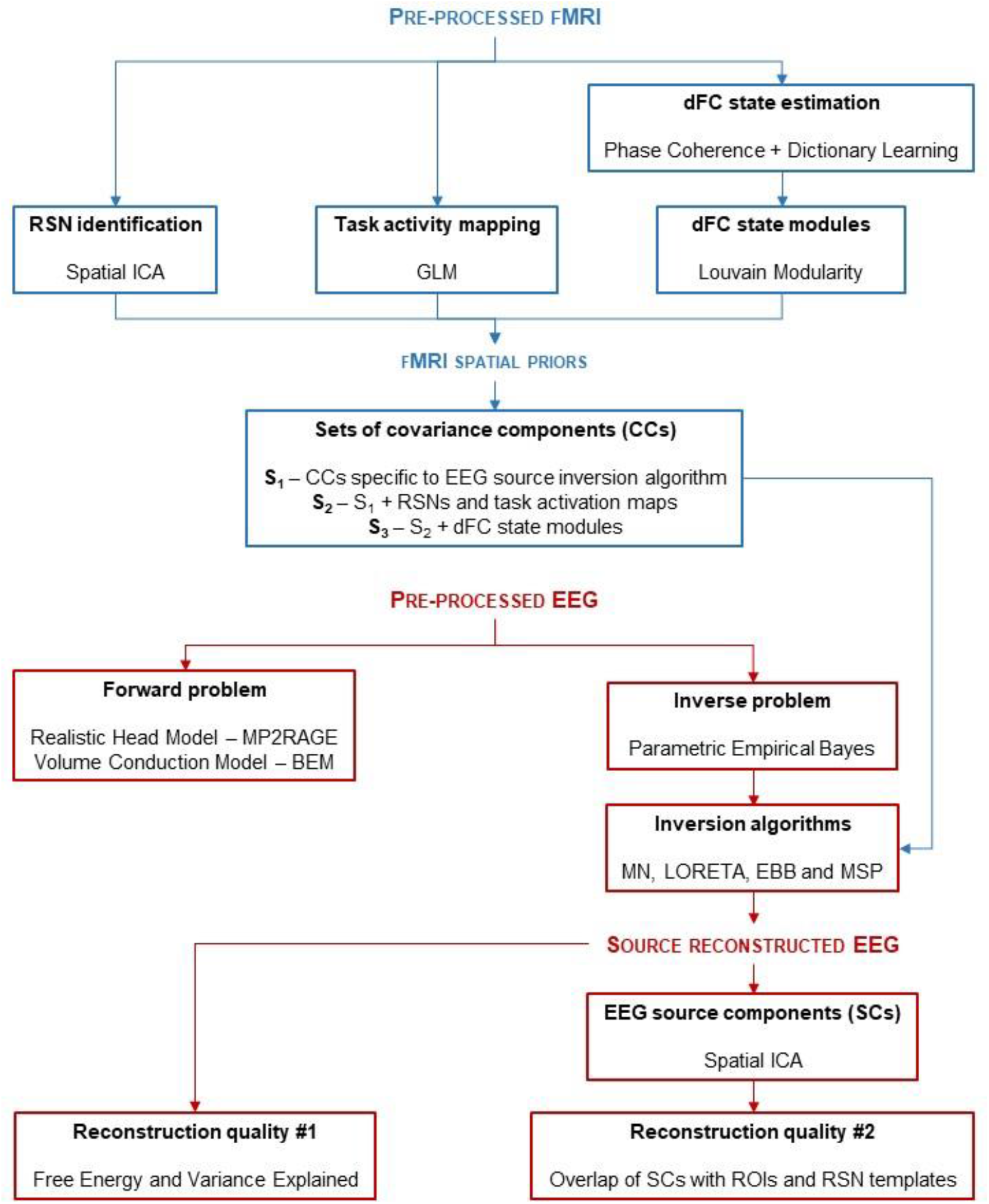
*Schematic diagram of the processing pipeline*. The pre-processed fMRI data is submitted to three different analyses in order to derive three types of fMRI spatial priors for EEG source reconstruction: 1) identification of RSNs through spatial ICA; 2) mapping of the task-related activity through GLM; and 3) by estimating the dFC fluctuations with phase coherence and the associated dFC states with dictionary learning, dFC state modules were obtained using the Louvain modularity algorithm. The covariance components (CCs) associated with these spatial priors were then included in several inversion algorithms, whose reconstruction quality was assessed by the free energy and variance explained of the associated models, and by the overlap of EEG source components (obtained through spatial ICA applied to the source reconstructed EEG) with ROIs and RSN templates.

#### 2.4.1 Pre-processing

The first 10 s of data were discarded to allow the signal to reach steady-state. Subsequently, slice timing and motion correction were performed using FSL tool MCFLIRT (Jenkinson et al., 2002), followed by a B_0_-unwarping step with FSL tool TOPUP (Andersson et al., 2003) using the reversed-phase encoding acquisition, to reduce EPI distortions. The distortion-corrected images were then corrected for the bias field using FSL tool FAST (Zhang et al., 2001), and non-brain tissue was removed using FSL tool BET (Smith, 2002). Nuisance fluctuations (including physiological noise) were then removed by linear regression using the following regressors (Abreu et al., 2017): 1) quasi-periodic BOLD fluctuations related to cardiac and respiratory cycles were modeled by a fourth order Fourier series using RETROICOR (Glover et al., 2000); 2) aperiodic BOLD fluctuations associated with changes in the heart rate as well as in the depth and rate of respiration were modeled by convolution with the respective impulse response functions (as described in Chang et al., 2009); 3) the average BOLD fluctuations measured in white matter (WM) and cerebrospinal fluid (CSF) masks (obtained as described below); 4) the six motion parameters (MPs) estimated by MCFLIRT; and 5) scan nulling regressors (motion scrubbing) associated with volumes acquired during periods of large head motion; these were determined using the FSL utility *fsl_motion_outliers*, whereby the DVARS metric proposed in (Power et al., 2012) is first computed, and then thresholded at the 75^th^ percentile plus 1.5 times the inter-quartile range. Finally, a high-pass temporal filtering with a cut-off period of 100 s was applied, and spatial smoothing using a Gaussian kernel with full width at half-maximum (FWHM) of 3 mm was performed.

For each subject, WM and CSF masks were obtained from the respective T_1_-weighted structural image by segmentation into gray matter, WM and CSF using FSL tool FAST (Zhang et al., 2001). The functional images were co-registered with the respective T_1_-weighted structural images using FSL tool FLIRT, and subsequently with the Montreal Neurological Institute (MNI) (Collins et al., 1994) template, using FSL tool FNIRT (Jenkinson et al., 2002; Jenkinson and Smith, 2001). Both WM and CSF masks were transformed into functional space and were then eroded using a 3 mm spherical kernel in order to minimize partial volume effects (Jo et al., 2010). Additionally, the eroded CSF mask was intersected with a mask of the large ventricles from the MNI space, following the rationale described in (Chang and Glover, 2009).

Each participant’s structural image was parceled into *N* = 90 non-overlapping regions of the cerebrum according to the automated anatomical labeling (AAL) atlas (Tzourio-Mazoyer et al., 2002). These parcels were co-registered to the participant’s functional space, and the pre-processed BOLD data were then averaged within each parcel.

#### 2.4.2 fMRI priors for EEG source reconstruction

From the pre-processed fMRI data, several potential priors for EEG source reconstruction were subsequently extracted (procedures described next), namely: resting-state networks for all runs, and task-related activity maps and dynamic functional connectivity (dFC) states for the task runs only.

##### Identification of resting-state networks

The pre-processed fMRI data were submitted to a group-level probabilistic spatial ICA (sICA) decomposition using the FSL tool MELODIC (Beckmann and Smith, 2004), whereby the data of each run for all participants is temporally concatenated prior to the sICA step, as recommended in the MELODIC’s guide for the identification of RSNs (https://fsl.fmrib.ox.ac.uk/fsl/fslwiki/MELODIC). The optimal number of independent components (ICs) was automatically estimated based on the eigenspectrum of its covariance matrix (Beckmann and Smith, 2004), with an average of approximately 40 ICs across runs.

An automatic procedure for the identification of well-known RSNs was then applied, in which the spatial maps of the ICs (thresholded at *Z* = 3.0) were compared with those of the 10 RSN templates described in (Smith et al., 2009), in terms of spatial overlap computed as the Dice coefficient (Dice, 1945). For each template, the IC map yielding the highest Dice coefficient was determined as the corresponding RSN. In the cases of non-mutually exclusive assignments, the optimal assignment was determined by randomizing the order of the RSN templates (a maximum of 10000 possible combinations were considered, for computational purposes), and then sequentially, and mutually exclusively, assigning them to the IC maps based on their Dice coefficient. The assignment with the highest average Dice coefficient across all RSN templates was then deemed optimal, yielding the final set of RSNs: three visual networks (RSN 1-3), the default mode network (DMN, RSN4), a cerebellum network (RSN5), a motor network (RSN6), an auditory network (RSN7), the salience network (RSN8), a right language network (RSN9) and a left language network (RSN10).

These 10 RSNs were then used as spatial priors for the reconstruction of sources of EEG collected on all four runs. RSNs were considered in the task runs because these have been shown to be also present in task-based studies (Cole et al., 2016; Di et al., 2013).

##### hMT+ and BM-related activity mapping

For the purpose of mapping hMT+/V5 from the localizer run, and the regions involved in the BM perception task from the other two runs, a general linear model (GLM) framework was used. For the localizer, two regressors representing the periods showing dots (stationary and moving) were considered. These regressors were built based on unit boxcar functions with ones during the respective periods, and zeros elsewhere. Similarly, three regressors representing the *body, foot* and *random* blocks of the BM runs were built for analyzing the BM runs, with the regressors also based on unit boxcar functions. All regressors were convolved with a canonical, double-gamma hemodynamic response function (HRF). The run-specific, HRF-convolved regressors were then included in a GLM that was subsequently fitted to the associated fMRI data using FSL tool FILM (Woolrich et al., 2001). The hMT+/V5 regions were identified from the localizer run by contrasting the moving and the stationary dots periods, whereas the areas associated with BM perception were mapped according to the following contrasts: *body* – *random, foot* – *random*, and *body* – *foot*. Voxels exhibiting significant changes within these contrasts were identified by cluster thresholding (voxel *Z* > 2.5, cluster *p* < 0.05).

In this way, a single spatial prior is obtained for reconstructing the sources of the EEG collected during the localizer run. In contrast, three spatial priors (one for each contrast) are made available for each of the two BM runs.

##### Dynamic functional connectivity analysis

The dFC analysis described here was only performed on the fMRI data collected during the task runs (localizer and two BM runs), since its purpose was to objectively identify a small set of dFC states associated with the tasks, and to use them as spatial priors in the subsequent reconstruction of EEG sources. The dFC was estimated using the phase coherence (PC) method, which allows to compute the dFC for each fMRI sample; the loss in temporal resolution and the ambiguous selection of a window size, both inevitable in conventional sliding-window correlation approaches (Preti et al., 2017), are thus avoided (Glerean et al., 2012). For the PC method, a second-order Butterworth band-pass filter in the range of 0.01–0.1 Hz was first applied to the parcel-averaged BOLD signals. The instantaneous phase, *θ*, of the filtered signal, *n*, was then estimated using the Hilbert transform (Cabral et al., 2017; Figueroa et al., 2019). For each participant, the dFC matrix **C** ∈ ℝ^*N*×*N*×*T*^ (*N* = 90 brain regions from the AAL atlas, and *T* is the number of fMRI samples, which depends on the run under analysis) was computed for each pair of parcels, *n* and *p*, and at each fMRI sample *t*, using the equation:

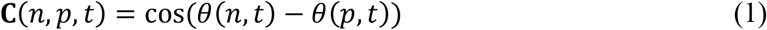

For each run and participant individually, the matrix **C** was then submitted to the leading eigenvector dynamics analysis (LEiDA) (Cabral et al., 2017; Figueroa et al., 2019; Lord et al., 2019), with the purpose of reducing the dimension of each temporal entry of **C** (*N*×*N*) by only considering the associated leading eigenvector (of dimension *N*), while nonetheless explaining most of the variance (> 50% in all cases, and up to 90%) (Lord et al., 2019). This step yielded the reduced dFC matrix **C**_**R**_ ∈ ℝ^*N*×*T*^, with the columns **c**_*t*_ ∈ ℝ^*N*×1^ (*t* = 1,…, *T*) representing the leading eigenvectors, and the rows indicating the parcels. Each eigenvector is composed by elements with positive and/or negative signs; if all positive, a global mode is governing the parcel-averaged BOLD signals where all the associated phases point in the same direction with respect to the orientation defined by the eigenvector (Figueroa et al., 2019). If the elements of the eigenvector have different signs, the parcels can be grouped into two networks according to their sign (positive or negative) in the eigenvector. The magnitude of the elements indicates how strongly the associated parcel belongs to its assigned network (Newman, 2006).

For the identification of dFC states, an *l*_1_-norm regularized dictionary learning (DL) approach was employed, following the methodology proposed in (Abreu et al., 2019). Briefly, this can be formulated as the matrix factorization problem **C**_**R**_ = **DA**, where **D** = [**d**_1_,…, **d**_*k*_] ∈ ℝ^*N*×*k*^ and **A** = [**a**_1_,…, **a**_*T*_] ∈ ℝ^*k*×*T*^ represent the dFC states and associated weight time-courses (i.e. contribution of each dFC state to reconstruct **C**_**R**_ at each time point), respectively; and *k* is the number of dFC states. These are estimated by solving the optimization problem given by:

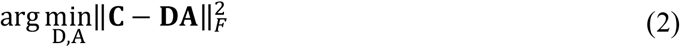

so that the reconstruction error of 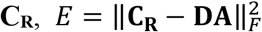, is minimized; ‖·‖_*F*_ denotes the Frobenius norm of a matrix. The estimation of **D** and **A** was performed using the algorithms implemented in the MATLAB® toolbox SPArse Modeling Software (SPAMS, Mairal et al., 2010). The sparsity of the solutions was controlled by a non-negative parameter λ on an *l*_1_-norm regularization framework. The number of dFC states *k* was varied between from 5 to 10 in unit steps, and λ between ten values from 1 to 0.1259 in decreasing exponential steps.

The optimal *k* and λ values were jointly determined with the dFC states to be considered as spatial priors in the EEG source reconstruction. This was achieved by first computing the Pearson correlation between the contrasts defined for identifying the activation maps (one for the localizer run, and three for the BM runs) and the dFC weight time-courses in **A**, for all possible combinations of *k* and λ. For the localizer run, the dFC state exhibiting the highest correlation across dFC states, and combinations of *k* and λ, was deemed as task-related. For the BM runs, the dFC state exhibiting the highest correlation across contrasts, dFC states and combinations of *k* and λ was first identified. For the optimal combination of *k* and λ, the most correlated dFC states associated with the remaining contrasts were then determined. In cases where multiple contrasts were associated with the same dFC state, only that state was considered for the subsequent analyses.

The dFC states of interest were then finally submitted to a modularity analysis, with the purpose of identifying modular (or community) structure in those states. Because the dFC states are column vectors, rather than square matrices representing a connectivity matrix as required for the modularity analysis, such connectivity matrix of a given dFC state **d**_*i*_ ∈ ℝ^*N*×1^ was first reconstructed by computing the outer product of the dFC state, 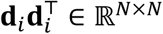 (Cabral et al., 2017). The Louvain algorithm as implemented in the Brain Connectivity Toolbox was then applied to the reconstructed connectivity matrices of the dFC states of interest (Rubinov and Sporns, 2010). This algorithm considers both the positive and negative weights of the unthresholded connectivity matrix, thus avoiding the ambiguous selection of a threshold as required in conventional modularity algorithms. Each of the *N* parcels is then assigned a label indicating which module the parcel belongs to. The network modules were then projected into binary 3D spatial maps to be used as spatial priors in the EEG source reconstruction, by identifying the voxels belonging to parcels (according to the AAL atlas used for parceling the brain) assigned to the same modules. The number of modules automatically identified by the Louvain algorithm was between 2 and 4 in all cases; in this way, the number of spatial priors built from this analysis varied according to the number of contrasts (run) and modules identified, with a maximum of 1 [contrast] × 4 [modules] = 4 for the localizer run, and 3 [contrasts] × 4 [modules] = 12 for the BM runs.

### 2.5 EEG data analysis

#### 2.5.1 Pre-processing

EEG data underwent gradient artifact correction on a volume-wise basis using a standard artifact template subtraction (AAS) approach (Allen et al., 2000) using the FMRIB tools implemented as a plug-in of the EEGLAB toolbox (Delorme and Makeig, 2004). The pulse artifact was removed using the method presented in (Abreu et al., 2016), whereby the EEG data is first decomposed using independent component analysis (ICA), followed by AAS to remove the artifact occurrences from the independent components (ICs) associated with the artifact. The corrected EEG data is then obtained by reconstructing the signal using the artifact-corrected ICs together with the original non-artifact-related ICs.

After the removal of the MR-induced artifacts, EEG data was then submitted to some of the routines of the automatic pipeline (APP) for EEG pre-processing described in (da Cruz et al., 2018), namely: 1) re-referencing to a robust estimate of the mean of all channels; 2) removal and interpolation of bad channels; and 3) removal of bad epochs of 1 second (matching the TR of the fMRI data). An additional ICA step was then performed with the purpose of removing additional sources of EEG artifacts; these were identified using the ICLabel algorithm (Pion-Tonachini et al., 2019), implemented as a plug-in of the EEGLAB toolbox (Delorme and Makeig, 2004). The classification provided by ICLabel is based on a previously trained model with a large EEG dataset collected outside the MR scanner, rendering this algorithm sub-optimal for our dataset. To cope with this, all ICs were visually inspected in order to validate, and eventually correct (for both false positives and negatives), the classification outputs of ICLabel. Finally, the EEG data was down-sampled to 500 Hz and band-pass filtered to 1–30 Hz.

#### 2.5.2 SSource reconstruction

The pre-processed EEG data from all runs was then submitted to several EEG source reconstruction procedures implemented in SPM12 (https://www.fil.ion.ucl.ac.uk/spm/). To reduce the computational load, the EEG data was further downsampled to a sampling rate of 60 Hz, two times the highest frequency component of the data.

##### The forward problem

A realistic head model was built by first segmenting each participant’s structural image into 3 tissue labels (brain, scalp and skull), and computing the deformation field needed to co-register the structural images into an MNI template. The individual meshes were then obtained by applying the inverse of this deformation field to the canonical meshes derived from the MNI template; meshes with 8196 vertices were considered. The electrode positions were co-registered to the scalp compartment by first considering their standard positions (in the 10/10 coordinate system), and then manually adjusting them to match the distortions clearly observed on the structural images. A realistically shaped volume conduction model was estimated using a boundary element model (BEM) with 3 layers (scalp, inner skull and outer skull). 8196 source dipoles were placed at the vertices of a cortical surface also derived from the MNI template and transformed into the structural image. The leadfield matrix was then estimated, mapping each possible dipole configuration onto a scalp potential distribution.

##### The inverse problem

The inverse problem was solved using a Parametric Empirical Bayesian (PEB) framework as implemented in SPM12, which can be formulated as (López et al., 2014):

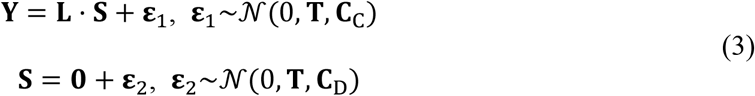

where **Y** ∈ ℝ^*C*×*T*^ is the EEG data with *C* channels (64 in this case) and *T* time samples (depends on the run under analysis); **L** ∈ ℝ^*C*×*D*^ is the leadfield matrix (*D* is the number of dipoles, 8196 in this case); and **S** ∈ ℝ^*D*×*T*^ is the unknown source dynamics at each dipole. 𝒩 (·) represents the multivariate Gaussian probability distribution, and **T** the known and fixed temporal correlations. The terms **ε**_1_ and **ε**_2_ denote the noise at the channel and source spaces, with covariance matrices **C**_C_ ∈ ℝ^*C*×*C*^ and **C**_D_ ∈ ℝ^*D*×*D*^, respectively. Channel noise is typically assumed to be uniform across channels, and therefore can be defined as **C**_C_ = *h*_C_**I**_C_, with *h*_C_ the channel noise variance and **I**_C_ ∈ ℝ^*C*×*C*^ the identity matrix. The source space covariance matrix **C**_D_ assumes the form:

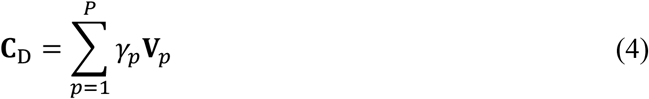

where **V**_*p*_ ∈ ℝ^*D*×*D*^ represents different types of covariance components (CCs) reflecting prior knowledge on the sources to be reconstructed, and *γ*_*p*_ the unknown hyperparameter denoting its relative importance. These hyperparameters work as regularization parameters in ill-posed problems such as the EEG inverse problem, and were estimated using a restricted maximum likelihood (ReML) algorithm that uses as cost function the free-energy of the model. Commonly used source inversion algorithms can then be derived from Eq. 3 by defining the CCs that appropriately reflect their assumptions. For instance, MN solutions assume that all dipoles have the same variance and no covariance; therefore, only one CC is defined as **V**_1_ = *h*_D_**I**_D_, with *h*_D_ the source noise variance and **I**_D_ ∈ ℝ^*D*×*D*^ the identity matrix.

In this work, we tested four source inversion algorithms: MN, LORETA, EBB and MSP; their derivations from Eq. 3 and associated CCs are thoroughly presented in (López et al., 2014). Additionally to those specific to a given algorithm, other CCs estimated from the fMRI-derived spatial priors (RSNs, activation maps and task-based dFC states) were considered (the procedures for their estimation are briefly described below). Specifically, all four inversion algorithms were tested using three different sets of CCs: the simplest set S_1_, with only CCs specific to the algorithm: 2) a larger set S_2_ comprising S_1_ and CCs from RSNs and activation maps (the latter for task runs only); and 3) the largest set S_3_ comprising S_2_ and CCs from the modules of the task-related dFC states (hence only tested on EEG collected from task runs). A total of 4 [inversion algorithms] × 3 [sets of CCs] = 12 reconstructions of EEG sources **S** were then performed for each subject and run (only 8 for the resting-state runs).

##### Estimation of covariance components from fMRI-derived spatial priors

CCs were estimated from the fMRI-derived spatial priors by first transforming them into binary priors. These 3D binary spatial priors were then projected onto the 2D cortical surface using nearest-neighbor interpolation (Henson et al., 2010), and smoothed using the Green’s function **G** of the cortical mesh adjacency matrix **M** ∈ ℝ^*D*×*D*^, **G** = *σ***M** (Harrison et al., 2007). The entries of **M**, *m*_ij_, are 1 if vertices *i* and *j* of the cortical surface are neighbors (within a defined radius) and 0 otherwise; here, a radius of 8 vertices and a smoothing parameter of *σ* = 0.6 were selected according to (Friston et al., 2008). The CCs of the smoothed (and projected) spatial priors are then obtained by computing their covariance matrices, i.e., their outer product. These procedures are illustrated in Fig. 2.

**Figure 2.**
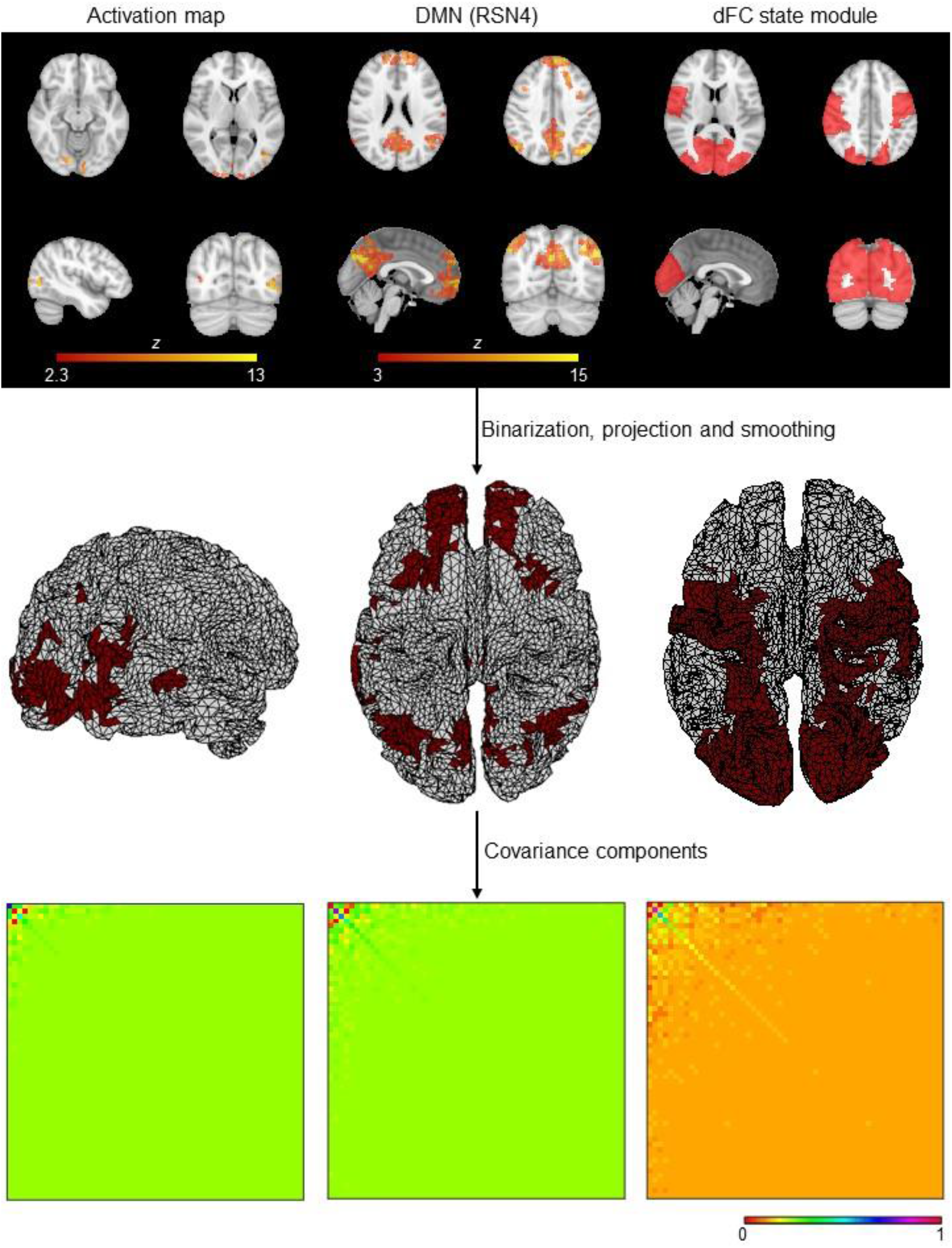
*Deriving covariance components (CCs) from fMRI spatial priors*. The 3D fMRI spatial priors are first binarized, projected onto the 2D cortical surface using nearest-neighbor interpolation and smoothed using the Green’s function. The associated CCs are then obtained by computing the outter product. For visualization purposes, the temporally reduced CCs are illustrated, by applying the same temporal projector considered when reducing the EEG data prior to its reconstruction.

### 2.6 Source reconstruction quality

#### 2.6.1 EEG source components

Following the rationale of previous studies (Abreu et al., 2020b; Liu et al., 2018, 2017), a spatial ICA step similar to that applied to the fMRI data for identifying RSNs was then performed on the reconstructed source dynamics **S**, with the purpose of separating those potentially associated with RSNs and/or other regions of interest in our tasks. This can be formulated as:

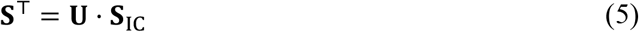

where **U** ∈ ℝ^*T*×*I*^ is the mixing matrix, with each column **u**_*i*_ ∈ ℝ^*T*×1^ the time-course of the source component (SC) *i*; and **S**_IC_ ∈ ℝ^*I*×*D*^ represents the spatial maps in the source space associated to each of the *I* SCs. Because the EEG data is submitted to a temporal reduction step prior to solving the inverse problem in order to reduce noise while guaranteeing a temporally continuous estimation of sources (López et al., 2014), the rank of **S** is reduced accordingly, being then defined an upper bound on the number of SCs to be estimated. Such maximum allowed number of SCs was then estimated, which was between 50 and 60 in all cases. Finally, the SCs were converted into *z*-scores, and the deformation field estimated while solving the forward problem was applied to transform them from the source space into the MNI space.

#### Quality metrics

Besides the free-energy (FE) of the inversion model and the variance explained (VE) of the reconstructed EEG data 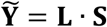 relative to the actual EEG data **Y** (see Eq. 3), other quality metrics reflecting more directly the presence of neuronal activity of interest in the SCs were considered.

First, because the perception of motion in general, and of biological motion in particular, is known to elicit certain brain regions, the following four spherical regions of interest (ROIs) of 10 mm centered at specific MNI coordinates (indicated in square brackets) were considered (Chang et al., 2018): anterior insula (aINS) at [±36, 24, 2], extrastriate body area (EBA) at [left –46, –75, –4; right 47, –71, –4], fusiform body area (FBA) at [left –38, –38, –27; right 43, –43, –28], and fusiform gyrus (FFG) at [±42, –56, –14]. Four additional task-related brain regions were obtained from FSL atlases (threshold applied to the probability maps is indicated in square brackets), namely (Chang et al., 2018): inferior frontal gyrus (IFG) [0.25], posterior superior temporal sulcus (pSTS) [0.25], visual area V3 [0.25] and visual area hMT+/V5 [0.10]. After binarizing the ROIs and the SC maps, the Dice coefficient *d*, and the proportion of the ROIs contained in the SC maps *p*_*RS*_, were then quantified according to (Dice, 1945):

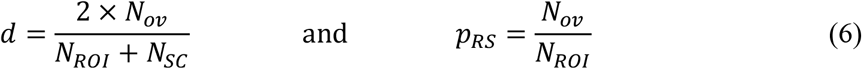

where *N*_*ROI*_ and *N*_*SC*_ denote the number of non-zero voxels in the ROIs and SC maps, respectively, and *N*_*ov*_ the number of overlapping non-zero voxels between the two images; both measures range from 0 (no overlap) to 1. These same two measures, *d* and *p*_*RS*_, were also computed between the SC maps and 10 RSN templates described in (Smith et al., 2009), in order to assess which, if any, SCs represented RSNs (similar to the identification of RSNs on fMRI data described previously).

All these measures were computed for each subject, run, inversion algorithm, set of covariance components (CCs), SC maps and maps of interest (8 ROIs and 10 RSN templates). Because only a subset of the SC maps is expected to be associated with those maps of interest, the SC map yielding the highest Gdice coefficient for each map was identified, and the associated *d*^*^ and 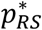 maximum values kept for subsequent analyses. The *d*^*^ and 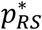 values were further summarized by computing their average within each map type (ROIs and RSN templates), thus yielding the final set of 13 [subjects] × 4 [runs] × 4 [inversions] × 3 [sets of CCs] × 2 [map types] = 1440 values of *d*^*^ and 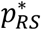.

#### 2.6.3 Statistical analysis

The main effects of the population group (MS patients and healthy subjects), inversion algorithm, the set of CCs and the type of map of interest, as well as interaction effects, were evaluated by means of a 4-way repeated measures Analysis of Variance (ANOVA) for the FE, VE, *d*^*^ and 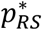 measures treated separately as the dependent variables. Multiple comparisons between the inversion algorithms, sets of CCs and interactions between the two were performed by means of a post-hoc statistical test with the Tukey-Kramer correction. A level of statistical significance *p<*0.05 was considered.

## 3 Results

In this work, the quality of EEG source reconstruction provided by the different combinations of (four) inversion algorithms and (three) sets of CCs, was first evaluated in terms of the FE and VE of the associated models, which are commonly considered in PEB frameworks. Because no significant differences were observed between population groups (healthy subjects and MS patients) the FE and VE values shown in Table 1 were averaged across participants; the values associated with the three visual perception task runs (hMT+/V5 functional localizer and two BM runs) were also averaged. The combination MSP+S_1_ (with S_1_ containing only CCs specifically associated with the inversion algorithm) yielded the lowest FE (the lower, the better) and the highest VE, only followed by LORETA+S_2_ and LORETA+S_3_. The ANOVA of the FE and VE values revealed significant main effects of the inversion algorithms and sets of CCs, as well as a significant interaction. The post-hoc tests on both FE and VE values showed no statistically significant differences between inversion algorithms, and the set S_1_ was significantly better than the sets S_2_ and S_3_; as expected, the combination MSP+S_1_ performed significantly better than other five combinations using the MN or EBB as inversion algorithms, and sets S_2_ or S_3_.

**Table 1:** *Average FE and VE values* across participants, and across three visual perception task runs, for all combinations of inversion algorithms and sets of covariance components. Values in bold represent the best across inversion algorithms for each CC set, and values in red represent the overall best (across inversion algorithms and CC sets).

| Sets of CCs | Inversion algorithms | Task runs (Localizer+BM) |  | Resting-state runs |  |
| --- | --- | --- | --- | --- | --- |
| | | $FE \times 10^4 (\pm \text{std})$ | $VE [\%] (\pm \text{std})$ | $FE \times 10^4 (\pm \text{std})$ | $VE [\%] (\pm \text{std})$ |
| $S_1$ | *MN* | 1.14±0.10 | 79.7±14.6 | 1.10±0.15 | 84.1±11.2 |
|  | *LORETA* | 1.13±0.10 | 79.5±14.5 | 1.10±0.15 | 83.9±11.3 |
|  | *EBB* | 1.13±0.11 | 79.4±14.6 | 1.09±0.15 | 83.2±11.8 |
|  | *MSP* | **1.10±0.11** | **84.5±10.5** | **1.06±0.15** | **87.6±8.3** |
| $S_2$ | *MN* | 1.13±0.11 | 81.8±13.7 | 1.09±0.15 | **85.6±10.6** |
|  | *LORETA* | **1.12±0.11** | **81.9±13.7** | **1.09±0.15** | 85.5±10.6 |
|  | *EBB* | 1.13±0.11 | 79.4±14.7 | 1.09±0.15 | 83.2±11.8 |
|  | *MSP* | 1.15±0.11 | 63.9±20.4 | 1.10±0.16 | 74.9±19.5 |
| $S_3$ | *MN* | 1.12±0.12 | 80.7±13.7 | NA | NA |
|  | *LORETA* | **1.12±0.12** | **80.7±13.7** | NA | NA |
|  | *EBB* | 1.13±0.12 | 78.2±14.8 | NA | NA |
|  | *MSP* | 1.16±0.12 | 60.3±15.5 | NA | NA |

In order to directly reflect the presence of neuronal activity of interest in the SCs, the source reconstruction quality was then quantified in terms of the overlap of SCs with the 8 ROIs and the 10 RSN templates. This is illustrated in Fig. 3, showing a considerable overlap (in terms of *d*^*^ and 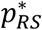) of two SCs with the EBA mask and the visual RSN1 template, for the first BM run of a given healthy subject. Consistently with the FE and VE values, the *d*^*^ and 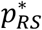 values were not statistically significantly different between population groups, and thus were averaged across participants and across task runs; these are depicted in Table 2. When considering only the CCs specific to the inversion algorithms (set S_1_), EBB yields the best results in all cases, and is the overall best (across sets of CCs) in terms of 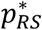 for the resting-state run. However, by combining S_1_ with RSNs and activation maps (set S_2_), MN achieves the highest *d*^*^ and 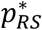 values for both types of runs, and the overall highest values (across sets of CCs) in terms of *d*^*^ for the resting-state run. For the task runs, the largest set of CCs including the dFC state modules (set S_3_) exhibits the overall best source reconstruction. Similarly to the statistical analysis of FE and VE, the ANOVA of the *d*^*^ and 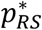 values revealed significant main effects of the inversion algorithms and sets of CCs, as well as a significant interaction. For the *d*^*^ values, the post-hoc statistical tests showed that MN and EBB inversion algorithms performed significantly better than LORETA and MSP, and that using the sets S_2_ or S_3_ was significantly better than only considering the set S_1_. The latter observation was also true for the 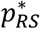 values, although in this case it was the MN and MSP inversion algorithms that yielded significantly better results than LORETA and EBB. The combinations EBB+S_2_ and MSP+S_3_ exhibited significantly higher *d*^*^ and 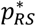 values, respectively, than a subset of combinations including the MN and LORETA algorithms, and the set S_1_.

**Figure 3.**
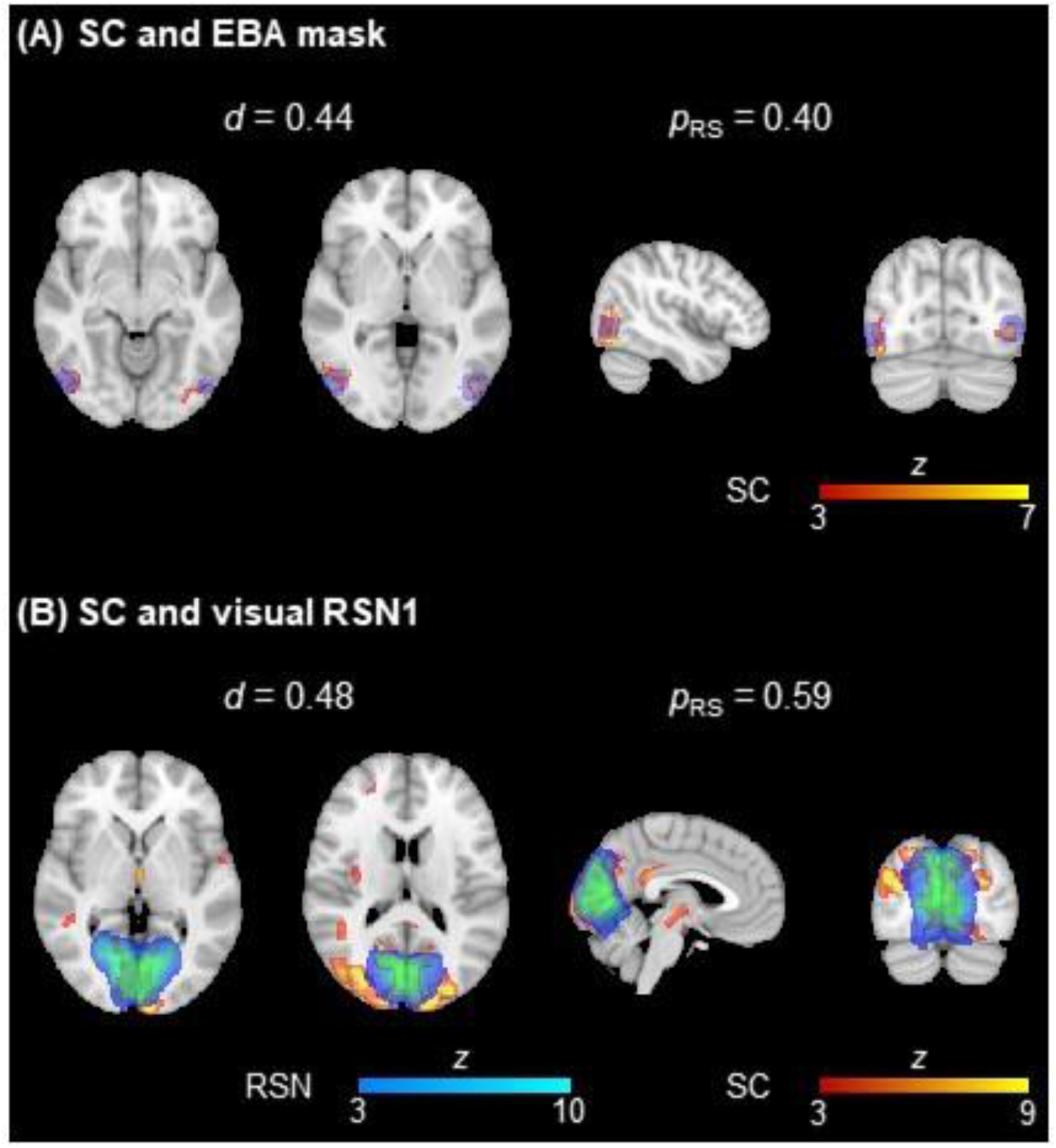
*Illustration of the overlap between two EEG SCs(in red-yellow) and (A) the EBA mask (in blue) and (B) a visual RSN (in blue-light blue)* from (Smith et al., 2009). The dice coefficient *d* and the proportion of the ROIs contained in the respective SCs are also depicted.

**Table 2:** *Average d*^*^ and 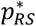 *values* across participants, and across three visual perception task runs, for all combinations of inversion algorithms and sets of covariance components. Values in bold represent the best across inversion algorithms for each CC set, and values in red represent the overall best (across inversion algorithms and CC sets).

| Sets of CCs | Inversion algorithms | Task runs (Localizer+BM) |  | Resting-state runs |  |
| --- | --- | --- | --- | --- | --- |
| | | $d^*$ (±std) | $p_{RS}^*$ (±std) | $d^*$ (±std) | $p_{RS}^*$ (±std) |
| $S_1$ | MN | 0.15±0.05 | 0.22±0.10 | 0.15±0.05 | 0.22±0.10 |
| | LORETA | 0.12±0.05 | 0.14±0.07 | 0.12±0.04 | 0.15±0.06 |
|  | EBB | **0.15±0.05** | **0.24±0.11** | **0.16±0.06** | **0.25±0.11** |
| | MSP | 0.14±0.05 | 0.18±0.09 | 0.12±0.04 | 0.16±0.09 |
| $S_2$ | MN | **0.15±0.05** | **0.24±0.12** | **0.16±0.05** | **0.24±0.11** |
| | LORETA | 0.15±0.05 | 0.23±0.12 | 0.15±0.04 | 0.22±0.12 |
| | EBB | 0.15±0.05 | 0.21±0.10 | 0.15±0.05 | 0.23±0.11 |
| | MSP | 0.14±0.05 | 0.23±0.11 | 0.14±0.05 | 0.22±0.09 |
| $S_3$ | MN | 0.15±0.05 | 0.24±0.12 | NA | NA |
| | LORETA | **0.16±0.05** | 0.24±0.12 | NA | NA |
| | EBB | 0.15±0.05 | 0.23±0.11 | NA | NA |
| | MSP | 0.12±0.07 | **0.35±0.10** | NA | NA |

## 4 Discussion

In this work, we aimed at optimizing the reconstruction of EEG sources by considering spatial priors derived from concurrently acquired fMRI data when solving the inverse problem, coupled with a systematic comparison of different inversion algorithms and sets of covariance components (CCs) reflecting those spatial priors, on a parametric empirical Bayesian (PEB) framework. The quality of the source reconstructions was quantified in terms of PEB-based metrics (the free-energy, FE; and the variance explained of the respective inversion models, VE), and physiologically-based metrics (the overlap of EEG source components with ROIs and RSN templates representative of brain activity of interest), the latter directly reflecting the presence of neuronal activity.

### EEG source reconstruction quality

Under a PEB framework, four inversion algorithms were tested here (MN, LORETA, EBB and MSP) for reconstructing sources from real EEG data collected from participants performing visual perception tasks and during rest, and considering three different sets of CCs. We found that depending on the type of quality metric (PEB- or physiologically-based), different conclusions could be taken. In terms of the PEB-based metrics (FE and VE), using the set consisting only of CCs specific to the inversion algorithms (S_1_) always yielded significantly better results than the CC sets including fMRI spatial priors (S_2_ and S_3_, comprising activation maps and RSNs, with or without dFC state modules, respectively). In contrast, by considering S_2_ and S_3_, the overlap of EEG source components (SCs) with the ROIs and RSN templates (measured by the dice coefficient *d*, and the proportion of the ROIs/RSNs contained in the SC maps, *p*_*RS*_) significantly surpassed that of S_1_. On the one hand, these contrasting results evidence the underlying optimization procedure used here, combining the PEB framework with the restricted maximum likelihood (ReML) algorithm for estimating the hyperparameters associated with each CC (Henson et al., 2010; Phillips et al., 2005). In fact, adding fMRI spatial priors drastically increases model complexity, which is penalized by ReML, and thus may explain the best PEB-based metrics when considering the more parsimonious inversion models (López et al., 2014). On the other hand, assessing the source reconstruction quality with metrics reflecting more directly the presence of neuronal activity of interest revealed that the information contained on the fMRI spatial priors is pivotal, suggesting that increasing model complexity in this way is needed for EEG SCs to contain such activity of interest, which is of the utmost interest for any subsequent analyses. Accordingly, the usefulness of fMRI spatial priors on EEG source reconstruction has already been shown in previous studies, with the addition of task-based activation maps (Henson et al., 2010; Lei et al., 2012, 2011, 2010) or RSNs (Lei, 2012) similar to those considered here improving the reconstructions. These however have only been compared in terms of conventional quality metrics, without taking explicitly into account the neuronal activity of interest. These observations highlight the relevance of using multiple quality metrics expressing different aspects of the reconstructed sources for more appropriately characterizing them, and consequently better informing the selection of the optimal reconstruction approach.

Regarding the optimal inversion algorithm, we found that no statistical differences were observed when comparing the FE and VE values, whereas MN and EBB yielded significantly higher *d* values than LORETA and MSP; MN and MSP achieved significantly higher *p*_*RS*_ values than EBB and LORETA. In contrast with the remaining inversion algorithms, LORETA is known for its low-resolution solutions (Michel and Brunet, 2019; Michel and Murray, 2012), which may render it inappropriate for localizing sources specifically associated with the limited number of rather small brain regions known to be involved in the tasks used in this study, and thus explaining its poorest performance (Halder et al., 2019). Concordant observations have been reported on previous comparison studies on simulated data (Bradley et al., 2016; Grova et al., 2006; Halder et al., 2019; Yao and Dewald, 2005), although no differences in performance were shown between MN and LORETA on real magnetoencephalography (MEG) or high-density EEG data (Hedrich et al., 2017). Similarly to MN, MSP has also been shown to provide solutions with high resolution (measured by the focal activation, for instance; Friston et al., 2008), despite potentially failing to fully recover the spatial extent of the sources (Grova et al., 2006). The same was observed for inversion algorithms of the family of beamformers as the EBB used here, namely the dynamic imaging of coherent sources (DICS) and linearly constrained minimum variance (LCMV), exhibiting higher focal activation and lower spatial extent than those of LORETA (Halder et al., 2019). Interestingly, our post-hoc interaction analyses showed that by combining EBB with S_2_ or MSP with S_3_, the best performance in terms of *d* and *p*_*RS*_, respectively, is achieved, suggesting that by coupling inversion algorithms designed for providing focal solutions with information derived from fMRI data, an optimal balance between specificity and sensitivity can be found. This recommendation is of particular relevance as it is drawn from the analysis of real EEG data (which contrasts with most studies in the literature that only focus on simulated data) that explicitly reflected the presence of neuronal activity in the reconstructed sources, rather than unspecific measures as those typically used in comparison studies.

Localizing in the brain the sources responsible for generating scalp EEG signals has been critical for determining the underpinnings of brain function in general, and those associated with multiple neurological disorders. Here, we compared for the first time EEG source reconstruction methods within a group of healthy subjects and MS patients, and found that the quality of the associated sources was irrespective of the presence of disease. This suggests that the recommendations made here may be extrapolated to future studies, regardless of the recruited population cohort.

### Reconstructing sources from EEG data collected simultaneously with fMRI

The accurate reconstruction of EEG sources is not only related with the appropriateness of the inversion models used, but also with the overall quality of the EEG signal (Liu et al., 2018). EEG data simultaneously acquired with fMRI is known to suffer from severe artifact contamination (Abreu et al., 2018), but state-of-the-art pre-processing pipelines as the one used here can now bring data quality to sufficiently high levels. Despite the potentially inevitable loss in data quality relative to EEG collected outside the MR scanner, the feasibility of reconstructing sources of EEG data acquired simultaneously with fMRI has already been demonstrated (Groening et al., 2009; Siniatchkin et al., 2010; Vulliemoz et al., 2010a, 2010b, 2009), particularly using EEG caps with a conventional spatial coverage (32 or 64 channels) as the one used in this study. Moreover, a direct relationship between EEG sources and fMRI networks has already been established first for data collected separately (Liu et al., 2017), and then validated on data collected simultaneously (Abreu et al., 2020b), supporting the feasibility of these procedures on this more challenging scenario. More importantly, analyzing EEG and fMRI data collected simultaneously is especially critical when studying spontaneous brain activity as the one associated with RSNs, for instance (Abreu et al., 2018). This further motivates the procedures performed here, and suggest that deriving spatial priors from fMRI data separately acquired from EEG data may be suboptimal, which in turn could scale down their potential for guiding the reconstruction of EEG. Future studies would need to be conducted to confirm this observation.

### Spatial priors and their relationship with EEG (sources)

In agreement with previous literature, in this work we found that including fMRI task activation maps and RSNs as additional CCs in the inversion models yielded significantly better EEG source reconstructions. This observation may be easily explained by the already known relationship between EEG and fMRI task activation and resting-state networks. In fact, source-reconstructed EEG data has already been used for mapping task-related fluctuations (Custo et al., 2014; Gonçalves et al., 2014), as well as for identifying RSNs (Abreu et al., 2020b; Liu et al., 2018, 2017), with recent studies showing a substantial overlap between these EEG maps and those typically obtained from fMRI data (Abreu et al., 2020b). Additionally, a relationship between fMRI RSNs and EEG has also been demonstrated in the sensor space, considering particularly the EEG rhythms extracted from the frequency domain (Goldman et al., 2002; Laufs et al., 2006; Moosmann et al., 2003; Scheeringa et al., 2008), which further supports the hypothesis that EEG carries in fact information that is also mapped with fMRI.

We then extended the exploration of fMRI spatial priors by also considering, for the first time, priors reflecting the fluctuations in the functional connectivity of task-related networks (dynamic functional connectivity, dFC). This was accomplished by estimating dFC fluctuations using phase coherence, followed by a dictionary learning step for finding the most recurrent dFC states, and a modularity analysis for identifying the network modules of the task-related dFC states. The rationale underlying our motivation for testing these spatial priors was based on recent literature showing that dFC fluctuations (Chang et al., 2013; Grooms et al., 2017; Korhonen et al., 2014; Omidvarnia et al., 2017; Preti et al., 2014; Tagliazucchi et al., 2012; Tagliazucchi and Laufs, 2015), and dFC states in particular (Abreu et al., 2020a; Allen et al., 2018), have distinct EEG correlates, which could also be reflected on source reconstructed EEG data. Our results evidence this because by adding the task-related dFC state modules as spatial priors, the quality of the source reconstruction further increased for the task runs, in terms of the overlap with the ROIs and RSN templates. Noteworthy, other than task-related dFC states were not considered here, because otherwise all dFC states would have to be included given the lack of criteria for selecting a subset of them, which would be necessary to control for the potentially increasing complexity of the models.

Importantly, spatial priors of different natures have already been suggested (Lei et al., 2015). Knowing the structural connectome by analyzing diffusion MRI (dMRI) data may inform functional connectivity measures in the EEG source space in terms of the strength of the underlying structural connections, by weighting those measures accordingly (Knösche et al., 2013). Moreover, the fiber tracking between regions of interest allows to estimate the time lag between their functional connections, which is of particular interest when considering distantly located regions (Chu et al., 2015). Effective functional connectivity estimates obtained through Granger causality in the EEG source space can also be informed by connectivity priors also derived from Granger causality analyses of the fMRI data, despite its much lower temporal resolution compared to that of EEG (Roebroeck et al., 2005). Dynamic causal modeling (DCM) also estimates effective functional connectivity by incorporating information at the meso-scale (described by neural models whose parameters are typically defined based on animal studies) and the macro-scale (Friston et al., 2019). The latter has parameters reflecting forward, backward and lateral connections between sources, which can be defined from dMRI and/or structural MRI data. Similarly to Granger causality, DCM can also be applied to fMRI data, and the results used as connectivity priors for EEG source reconstruction (Lei et al., 2015). Naturally, these connectivity priors may be more crucial for studying EEG functional connectivity in the source space, which was not the case in the present study.

An alternative to the PEB framework for incorporating priors is the use of penalty functions (Lei et al., 2015). These constraint the inverse solutions using different types of norms and weight matrices that indirectly reflect a given prior, from which the MN and LORETA algorithms used here can be defined. Penalty functions have the advantage of easily balancing between sparse and smooth solutions by simply adjusting the norm accordingly, or combining multiple terms with different norms for intermediate solutions (Valdés-Sosa et al., 2009). However, the ability of explicitly incorporating spatial priors as covariance components in the inversion models designed under a PEB framework render it more interpretable, and therefore potentially more suitable for testing different types of fMRI spatial priors as it was performed here (López et al., 2014). Regardless, extending the present study by also comparing these two frameworks would be interesting to further inform researchers on not only the best combination of inversion algorithms and sets of covariance components, but also on the optimal framework.

## Conclusions

In this study, we systematically compared the quality of the source reconstruction of EEG data performed using different combinations of four inversion algorithms and three sets of covariance components incorporating different types of spatial priors derived from concurrently acquired fMRI data. We found that according to the quality metrics reflecting the presence of neuronal activity, combining the EBB or MSP algorithms with CC sets including fMRI task activation maps and RSNs yields the overall best source reconstruction, and that by further including dFC state modules as spatial priors, the quality of EEG sources from the task runs is optimal. We show that incorporating fMRI spatial priors in general, and for the first time dFC state modules in particular, is thus crucial for optimizing the reconstruction of EEG sources (and consequently any subsequent analyses). By providing a clear recommendation on the best approach for tackling the challenging inverse problem supported by our comprehensive analyses, we believe that future studies, particularly using real EEG data, may then be more informatively guided on this intricate research field.

## Acknowledgements

This work was supported by Grants Funded by Fundação para a Ciência e Tecnologia, PAC –286 MEDPERSYST, POCI-01-0145-FEDER-016428, BIGDATIMAGE, CENTRO-01-0145-FEDER-000016 financed by Centro 2020 FEDER, COMPETE, FCT UID/4950/2020 – COMPETE, CONNECT.BCI POCI-01-0145-FEDER-30852, and BIOMUSCLE PTDC/MEC-NEU/31973/2017. FCT also funded an individual grant to JVD (Individual Scientific Employment Stimulus 2017 - CEECIND/00581/2017).

## Notes

### Competing Interest Statement

The authors have declared no competing interest.

